# Fitness trade-off and the discovery of a novel missense mutation in the PmrB sensor kinase of a colistin-resistant *Pseudomonas aeruginosa* strain developed by adaptive laboratory evolution

**DOI:** 10.1101/2024.10.31.621426

**Authors:** Indira Padhy, Sambit K. Dwibedy, Saswat S. Mohapatra

**Author notes:** For correspondence: **Saswat S. Mohapatra**, Ph.D., Assistant Professor, Department of Biotechnology, Berhampur University, Bhanja Bihar, Berhampur- 760007.

## Abstract

*Pseudomonas aeruginosa* is a prominent bacterial pathogen that causes several nosocomial infections and is notorious for its environmental resilience and rapid development of resistance to frontline antibiotics. A major cause of mortality and morbidity among cystic fibrosis patients, multidrug-resistant *P. aeruginosa* is often targeted with the antibiotic colistin as a last option. However, increasing reports of colistin resistance among *P. aeruginosa* is a matter of significant concern. Though the molecular mechanisms responsible for the development of colistin resistance are well known, the evolutionary trajectory to colistin resistance is an important area of investigation. In this work, using the adaptive laboratory evolution (ALE) approach we have evolved a colistin-sensitive *P. aeruginosa* ancestral strain to a resistant one. During the process of laboratory evolution in 106 generations, colistin MIC was increased 32-fold. The evolved strain had lower fitness than the ancestral strain as evidenced by a lower growth rate and higher doubling time. Moreover, the evolved strain produced more biofilm and less pyocyanin. Interestingly, the evolved strain showed collateral sensitivity to several antibiotics such as co-trimoxazole, rifampicin, kanamycin, tigecycline, penicillin, ampicillin, and teicoplanin. On analysing various TCS modules involved in the development of colistin resistance a novel missense mutation (V136G) was detected in the PmrB sensor kinase. Bioinformatics prediction indicated that the mutation could be deleterious, though the functionality of the PmrB mutant remains to be validated experimentally.

## Introduction

Antimicrobial resistance (AMR) has been among the most significant challenges in the healthcare sector in the last few decades. AMR’s global impact has earned it the reputation of being a “silent pandemic”. As per an estimate 4.95 million deaths globally were associated with antibiotic-resistant bacteria in the year 2019 alone (Murray et al., 2022), which is predicted to rise to more than 8.22 million in the year 2050 (Naghavi et al., 2024). Among the bacterial species significantly associated with the rise of drug-resistant infections, ESCAPE pathogens (for *Enterococcus faecium, Staphylococcus aureus, Klebsiella pneumoniae, Acinetobacter baumannii, Pseudomonas aeruginosa*, and *Enterobacter*) are most important, against whom urgent antimicrobial intervention methods are warranted (Kelly et al., 2024). Among the ESCAPE pathogens, *P. aeruginosa* is notorious for causing a wide range of life-threatening acute and chronic infections like respiratory infections, urinary tract infections, bacteraemia etc. specifically among the immunocompromised patients (Laborda et al., 2021; Weimann et al., 2024). *P. aeruginosa* is also the major causative agent of morbidity and mortality in cystic fibrosis patients. Moreover, *P. aeruginosa* shows remarkable adaptability in various adverse environmental conditions and is known for acquiring resistance to a variety of frontline antibiotics in a short period making the infections difficult to treat (Laborda et al., 2021; Rossi et al., 2021; Weimann et al., 2024). Multidrug-resistant (MDR) *P. aeruginosa* is often treated with colistin, a lipopeptide antibiotic of the polymyxin class, which remains the drug of last choice (Padhy et al., 2024)

With the increasing use of colistin to treat *P. aeruginosa* infections, resistance to colistin is emerging in the clinic. Recently, in India, the rate of colistin resistance among the *P. aeruginosa* strains isolated from nosocomial infections was found to be 13.3% (Dwibedy et al., 2024). Several mechanisms responsible for colistin resistance in *P. aeruginosa* have been identified, of which LPS modification with L-Ara4-N and pEtN encoded by the *arnBCADTEF*-*ugd* and *eptA* operons respectively are the most significant (Mohapatra et al., 2021; Padhy et al., 2024). This modification of the LPS results in decreased interaction with the colistin, and is mediated via two-component signal transduction systems (TCS) (Padhy et al., 2024). To date, at least five TCSs namely PhoPQ, PmrAB, ParRS, CprRS, and ColRS, have been identified to play a role in developing colistin resistance in *P. aeruginosa*, of which PhoPQ and PmrAB are the most well-characterized (Jochumsen et al., 2016; Dößelmann et al., 2017; Padhy et al., 2024).

With the increasing reports of colistin-sensitive *P. aeruginosa* becoming resistant to it in clinical settings, it is important to understand the mechanisms of the emergence of resistance in response to drug exposure (Dößelmann et al., 2017; Wüllner et al., 2022; Antunes et al., 2024). Adaptive laboratory evolution (ALE) or experimental evolution has become an important tool for understanding the evolutionary trajectory of various bacterial species to antibiotics and other environmental conditions (Lenski, 2017; Cooper, 2018; Van Den Bergh et al., 2018). ALE involves the continuous growth of the bacteria in increasing concentrations of the drug *in vitro* and comparing the ancestral and evolved bacteria for the development of antibiotic resistance and other associated phenotypes (Van Den Bergh et al., 2018; Antunes et al., 2024). To comprehend the evolutionary mechanisms, *P. aeruginosa* strains have been previously tested against several antibiotics both *in vivo* and *in vitro* by using ALE approach (Jochumsen et al., 2016; Dößelmann et al., 2017; Laborda et al., 2022; Disney-McKeethen et al., 2023; Filipow et al., 2023; Antunes et al., 2024; Higazy et al., 2024; Weimann et al., 2024). These studies have provided crucial insights regarding genes and pathways primarily impacted during the adaptation process and their role in natural selection.

In the present study, we have used ALE to evolve a highly colistin-resistant strain of *P. aeruginosa* by serially propagating the ancestral sensitive strain in gradually increasing concentrations of colistin for 16 days to obtain a colistin-resistant strain that can grow at 130 µg/ml of colistin. While developing the colistin resistance the evolved strain was found to have lost its fitness as evident from various growth assays. Moreover, we have detected a novel missense mutation in the PmrB sensor kinase (V136G) which might have a role in the development of colistin resistance.

## Materials and Methods

### Bacterial strain

The *P. aeruginosa* strain belonging to the PAO group used in the study was a kind gift from Dr. Harapriya Mohapatra, NISER, Bhubaneswar, India. The strain was routinely grown in LB medium and Mueller-Hinton broth as and when required.

### Adaptive laboratory evolution (ALE) experiment

The ALE experiment was used to develop a colistin-resistant *P. aeruginosa* strain by growing it in increasing concentrations of colistin following the protocol described earlier (Vinchhi et al., 2023), with few modifications. The overall scheme of the evolution experiment is presented in Fig. 1 and as follows. The colistin-sensitive *P. aeruginosa* strain designated PA (An) was retrieved from the freezer and cultured on an LB agar plate. A single colony from the plate was inoculated in a test tube containing 3 ml of cation-adjusted Mueller-Hinton broth (CAMHB) and incubated at 37°C and 150 rpm overnight. The overnight starter culture was diluted 100-fold in a fresh MH broth (30 ml) supplemented with 1 µg/ml of colistin and incubated at 37°C with 150 rpm for 24 hours to allow the cells to adapt. Subsequently, the experimental evolution was continued with daily transfer (after 24 hrs.) of 1% culture to a fresh MHB medium containing a higher colistin amount. The colistin concentration was gradually increased in the medium to allow the bacteria sufficient adaptation time and limit population extinction. The growth conditions remained constant for the entire experimental evolution period. After a total of 16 transfers (approx. 106 generations), a *P. aeruginosa* strain growing at 130 µg/ml of colistin was obtained and designated as PA (Evo). The strain was maintained on an LB agar plate with 130 μg/ml colistin selection for further analysis. During the experiment, the *P. aeruginosa* strain was also grown without antibiotic selection as an internal control.

**Fig. 1.**
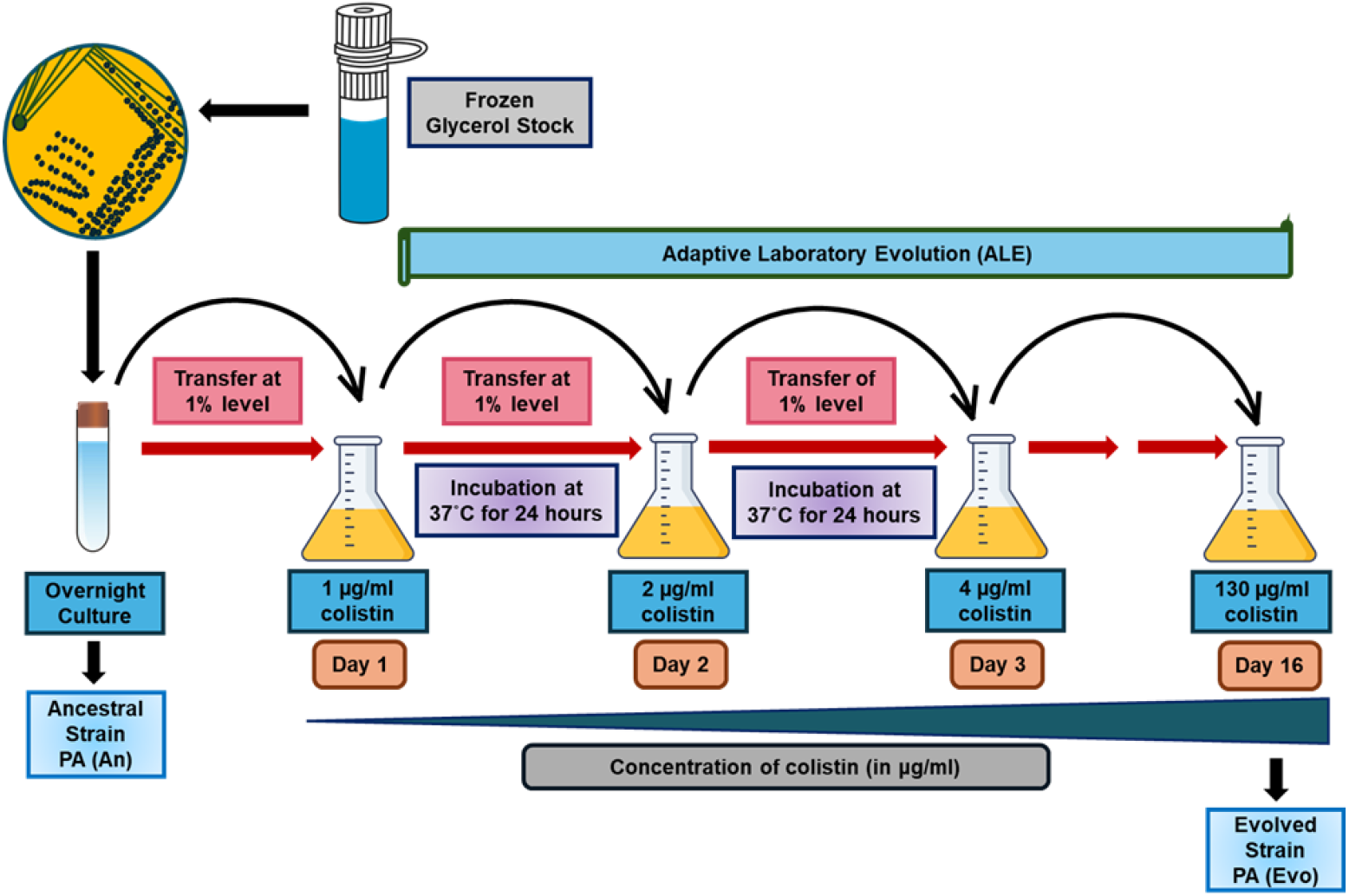
The scheme of the adaptive laboratory evolution (ALE) experiment followed in the current study. PA (An) strain was retrieved from the frozen stock and grown on LB agar followed by transferring a colony to MH broth and growing overnight. Adaptation to colistin started with transferring 1% inoculum to a fresh MH broth with 1 µg/ml for 24 hrs. The serial passage of inoculum to increasing concentration of colistin was continued for 16 days (106 generations) to obtain a strain PA (Evo) that could grow on 130 µg/ml of colistin.

### Growth characteristics

The colony characteristics of the PA (An) and the PA (Evo) strains were compared by growing them on LB agar plates. To assess their growth in the liquid medium, the strains were grown overnight in LB broth, diluted 1:100 in fresh LB medium, and grown till the stationary phase. The optical density at 600 nm (OD_600_) was measured each hour until 12 hrs. A growth curve was plotted using the OD_600_ vs time using the Microsoft Excel program. From the exponential part of the growth curve the growth rate (µ) and the doubling time (Td) were estimated from the formula below,

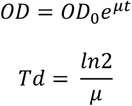

Where OD_0_ is the initial optical density, OD is the optical density at time t, and e is the exponential function. The colony counts were also determined by plating a suitably diluted culture (100 µl) on an LB agar plate. The log_10_ (CFU/ml) vs time was plotted using the Microsoft Excel program. These experiments were performed in triplicate and the mean value and the standard deviation were used for the calculations.

### Minimum inhibitory concentration (MIC) determination

The MIC for colistin against the PA (An) and the laboratory-evolved PA (Evo) strains was determined using the E-test and the broth microdilution methods (Joyce et al., 1992; CLSI, 2018). In the case of the E-test, the strains grown up to 0.5 McFarland standard in MHB were spread over the MH agar and the colistin EZY MIC strip (EM020, Himedia, Mumbai) was placed on it. The plate was incubated for 12-16 hours, and the result was interpreted by observing the point of intersection of the ellipse (Joyce et al., 1992).

The MIC determination using the broth microdilution method was performed following the protocol described earlier (CLSI, 2018; Satlin et al., 2020). Briefly, the colistin concentration range was prepared by a 2-fold serial dilution method. Meanwhile, the overnight grown culture was diluted 1:100 in MHB media and incubated in a shaking condition till the turbidity of the suspension reached 0.5 McFarland standards, which corresponds to an optical density containing 1.5×10^8^ CFU/ml. This suspension was again diluted 150 times using fresh MHB resulting in an inoculum of 10^6^ CFU/ml. This diluted culture was added into each antibiotic dilution well in a 96-well plate to bring the final inoculum to 5×10^5^ CFU/ml. The experiment was performed in triplicate. The 96-well plate was incubated at 37°C for 20 hrs. The well containing the lowest concentration of antibiotics, with no visible growth, was considered the MIC.

### Antibiotic susceptibility assay

Antimicrobial susceptibility testing of the strains PA (An) and PA (Evo) was performed using the Kirby-Bauer disk diffusion method (CLSI, 2020). Antibiotics with different modes of action belonging to different classes were obtained from Himedia, Mumbai. The antibiotics tested were Carbapenem [ertapenem (10 mcg), imipenem (10 mcg) and meropenem (10 mcg)]; Glycopeptide [vancomycin (30 mcg) and teicoplanin (30 mcg)]; Cephalosporin [cefotaxime (30 mcg) and cefepime (30 mcg)]; Macrolide [erythromycin (15 mcg) and azithromycin (15 mcg)]; Penicillin [methicillin (5 mcg), ampicillin (10 mcg), piperacillin (100 mcg) and penicillin-G (10 mcg)]; Tetracycline [tetracycline (30 mcg), doxycycline (30 mcg) and tigecycline (15 mcg)]; Aminoglycoside [streptomycin (300 mcg), gentamycin (120 mcg), tobramycin (10 mcg), amikacin (30 mcg) and kanamycin (30 mcg)]; Quinolone [moxifloxacin (5 mcg), lomefloxacin (10 mcg), ofloxacin (5 mcg), norfloxacin (10 mcg), ciprofloxacin (5 mcg), and nalidixic acid (30 mcg)]; Sulphonamide [trimethoprim (5 mcg) and co-trimoxazole (25 mcg)]; Rifampicin (5 mcg); Chloramphenicol [chloramphenicol (30 mcg)]; Lincosamide [clindamycin (2 mcg)]; Monobactam [aztreonam (30 mcg)] and Nitrofurantoin [nitrofurantoin (300 mcg)]. The zone of inhibition was measured using an antibiotic zone scale. Susceptibility results were interpreted following the Clinical Laboratory Safety Institute guidelines (CLSI, 2020).

### Biofilm assay

The biofilm assay was performed as described earlier (Merritt et al., 2006). Briefly, the strains PA (An) and PA (Evo) were incubated overnight in tryptic soya broth (TSB) supplemented with 4% glucose. Fresh growth started in enriched TSB by transferring 1% of the starter culture and from each test tube, 250 µl of the sample was dispensed in 4 wells of a microtiter plate. The plate was covered and incubated for 24 hours at 37°C. After incubation, the planktonic cells were discarded leaving the biofilm intact. The biofilm was washed with 250 µl of 0.8% NaCl, stained with 0.1% (w/v) crystal violet and incubated at room temperature for 45 minutes. Then the plate was blotted vigorously on a stack of paper towels to remove excess dye and dried overnight in an inverted position. To each well 250 µl of 30% acetic acid was added, mixed by pipetting and retained for 15 minutes. The absorbance of each well was measured in an i-mark microplate reader (Biorad, USA) at 595 nm using 30% acetic acid plus TSB stained with CV without culture as blank. The experiment was performed in triplicate.

### Pyocyanin production assay

Pyocyanin assay was performed to determine the amount of pyocyanin produced by the strains using the procedure described earlier (Zhu et al., 2019), with few modifications. Briefly, overnight cultures were adjusted to OD_600_ 1.0 before inoculating in 25 ml of LB with 1:100 dilution and incubated for 24 hours at 37°C, 200 rpm in 250 ml flasks. The cultures were centrifuged at 8000 g for 10 min, and the supernatant was collected. Supernatants (7.5 ml) were transferred to three centrifuge tubes of 15 ml volume to which 4.5 ml of chloroform was added and vortexed for 10-20 seconds till the colour became greenish blue. The mixture was centrifuged for 10 min at 8000 g. The resulting blue layer (approx. 3 ml) at the bottom consisting of chloroform and pyocyanin was transferred to a new tube. To the mixture, 1.5 ml of 0.2 M HCl was added and vortexed for 10-20 seconds which resulted in the conversion of blue colour into pink. Samples were centrifuged for 2 min at 8000 g, and 1 ml from the upper pink layer was used to estimate the pyocyanin concentration by checking the absorbance at 520 nm. This experiment was performed in triplicate. Pyocyanin concentration (μg/ml) was calculated by multiplying the obtained A_520_ value with 17.072.

### Genomic DNA isolation and PCR

The genomic DNA of the PA (An) and PA (Evo) strains were prepared following the procedure described previously (Wilson, 2001). The quality and quantity of the genomic DNA preparation were assessed using agarose gel electrophoresis and measuring the absorbance at 260 nm and 280 nm (A_260_/A_280_).

The genomic DNA was used for the amplification of histidine kinase and response regulator genes belonging to the PhoPQ and PmrAB two-component signal transduction systems (TCSs) using the primers listed in Table 1. Briefly, the Emerald Amp GT PCR Master Mix (DSS Takara, India) was used to amplify the genes *phoQ, phoP, pmrB*, and *pmrA* following the manufacturer’s instructions. The following cyclic conditions were used for the amplification of the genes; initial denaturation at 94°C for 5 min, denaturation at 94°C for 30 sec, annealing at 57-59°C for 30 sec, extension at 72°C for 45 sec, for 35 cycles, followed by final extension at 72°C for 10 min. The PCR products were checked by agarose gel electrophoresis. The amplicons were purified using the HiPurA® PCR product purification kit (Himedia, Mumbai).

**Table 1.**
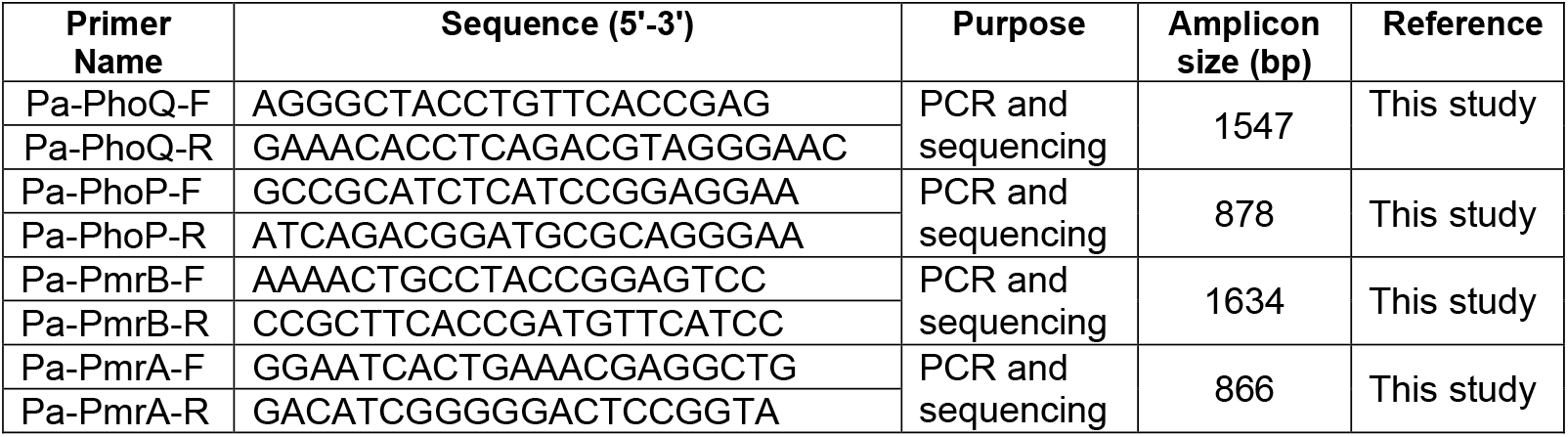
List of primers used in the study.

### DNA sequencing and analysis

The *phoQ, phoP, pmrB*, and *pmrA* amplicons were sequenced at the Barcode Biosciences, Bangalore using the Sanger sequencing method. The nucleotide sequences were analysed using the MEGA program version 11.0 (Tamura et al., 2021). Multiple sequence alignment using the gene sequences from the PA (An) and PA (Evo) strains was performed using the ClustalW. The nucleotide sequences have been submitted to the NCBI GenBank with the accession numbers PQ014646 to PQ014653.

Based on the nucleotide sequence information, the non-synonymous mutation was detected. The impact of the novel non-synonymous substitution mutation in PmrB (V136G) sensor kinase was analysed using the PROVEAN (Protein Variation Effect Analyzer) webserver (Choi and Chan, 2015) using a default threshold value of -2.5.

## Results

### Growth characteristics of the evolved strain

In this study, to understand the mechanisms of emergence of colistin resistance in *P. aeruginosa*, we employed the ALE approach to produce a highly resistant strain (Fig. 1). ALE was performed by growing the PA (An) strain in the Muller Hinton Broth (MHB) with increasing concentration of colistin through a serial passaging for 16 days (106 generations) to obtain the evolved PA (Evo) strain that can grow in presence of 130 μg/ml of colistin. The PA (Evo) strain was compared with PA (An) on growth and other phenotypic characteristics.

On growing the strains on LB agar without antibiotic selection, the PA (Evo) colonies were found to be smaller in comparison to the PA (An) (Fig. 2A). Moreover, they did not produce the characteristic green pigment generally associated with *P. aeruginosa*. The colonies took longer to appear on the plate indicating a slower growth rate or a longer generation time (Fig. 2A). To further compare their growth characteristics, the strains were cultured in LB broth without colistin selection, and the growth was followed for 12 hours (Fig. 2B). As evident from the growth curve obtained with OD_600_ measurement, the evolved strain grew considerably slowly, and the onset of the log phase was delayed by more than 1 hr (Fig. 2B). Further, the final growth yield was lesser in PA (Evo) as compared to the PA (An) strain; as the former reached the stationary phase at OD_600_ of around 1.5, whereas the latter reached at OD_600_ of 2. Similarly, when the log_10_ (CFU/ml) was plotted against time for the strains, the PA (An) was found to reach exponential growth one hour before the PA (Evo) strain, though the final number of viable cells were of a similar order of magnitude (Fig. 2C).

**Fig. 2.**
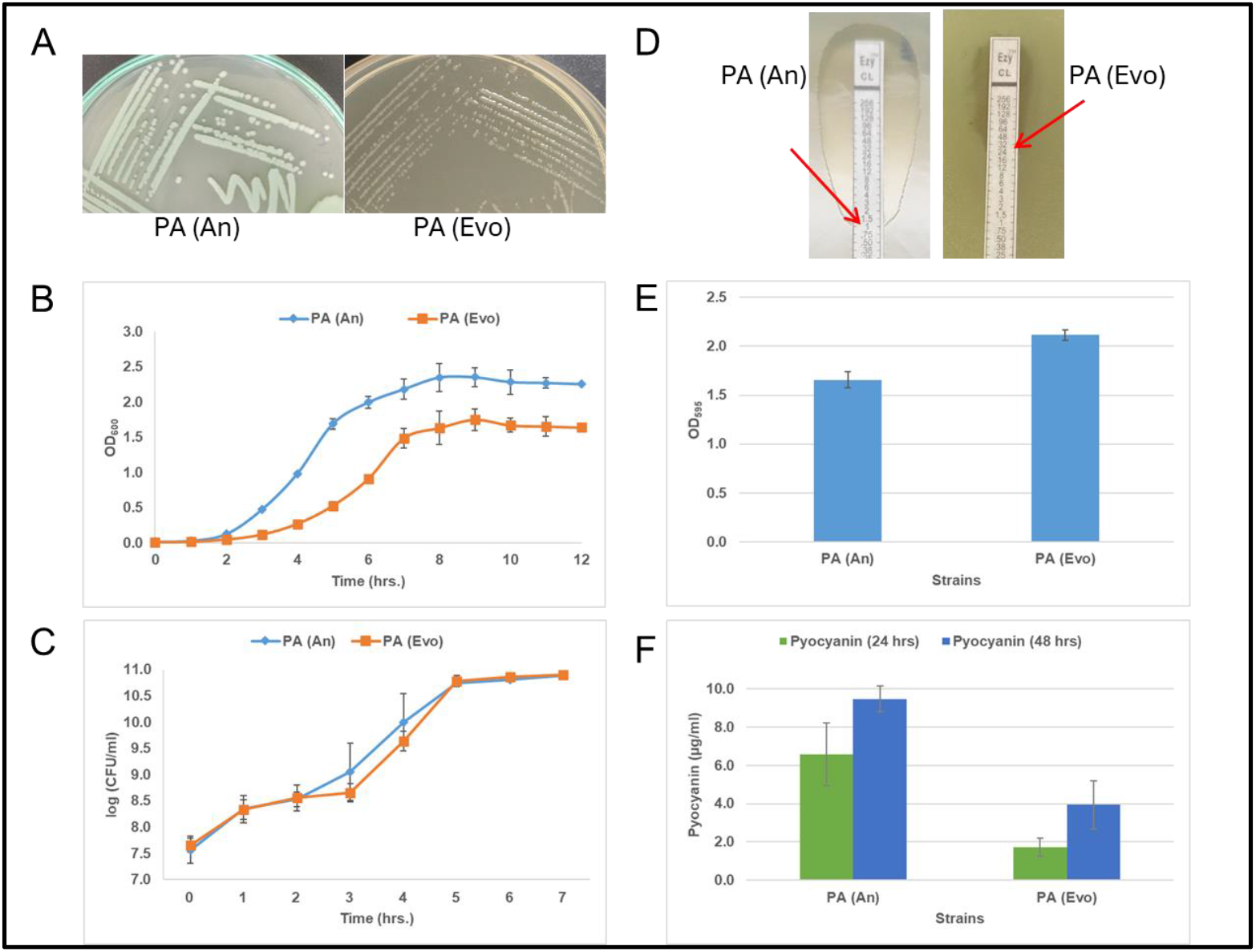
Phenotypic characterization of the PA (Evo) strain and comparison with the ancestral PA (An) strain. **(A)** Growth on LB agar plates, **(B)** Comparative growth analysis in LB broth plotting the OD_600_ vs time, **(C)** Comparative growth analysis in LB broth plotting the log_10_ (CFU/ml) vs time, **(D)** Determination of colistin MIC using the E-test. The red arrow indicates the colistin MIC value, **(E)** Comparison of biofilm formation between the PA (An) and PA (Evo) strains, and **(F)** Comparison of pyocyanin production by the strains at 24 hrs and 48 hrs.

The growth rate (µ) and the doubling time (Td) were estimated from the growth assay as described in the materials and methods. When compared, the growth rate of the PA (An) strain was 1.188 hr^-1^ and the same for the PA (Evo) was 0.858 hr^-1^ (Table 2). While the doubling time for the PA (An) was 35 min in the LB broth, it was 42.16 min for the PA (Evo) strain (Table 2), clearly indicating that the strain has a reduced fitness while acquiring the colistin resistance phenotype.

**Table 2.**
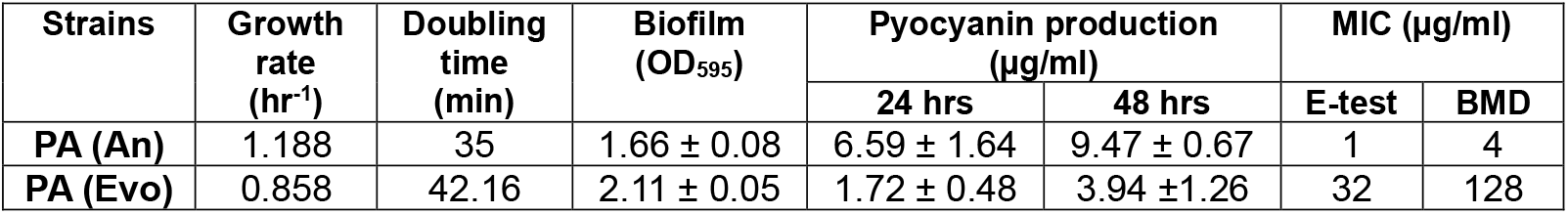
Comparative growth and other phenotypic characteristics of the PA (An) and PA (Evo) strains.

### Antibiotic susceptibility profiling and detection of collateral sensitivity

The colistin MIC was determined for the PA (An) and the PA (Evo) strains using the E-test and the broth microdilution (BMD) method. While the colistin MIC of the PA (An) was 1 and 4 μg/ml respectively as determined using the E-test and the BMD, the MIC for the PA (Evo) was found to be 32 and 128 μg/ml using the same methods respectively (Table 2, Fig. 2D). The comparison indicates that there is at least a 32-fold increase in the colistin MIC during the evolution process.

The PA (An) and the PA (Evo) strains were checked for their antibiotic susceptibility pattern using the Kirby-Bauer disc diffusion method. While acquiring the colistin resistance the PA (Evo) strain interestingly showed collateral sensitivity to several antibiotics tested, particularly towards penicillin G, ampicillin, teicoplanin, streptomycin, kanamycin, co-trimoxazole, and rifampicin (Table 3).

**Table 3.** Comparative antibiotic susceptibility profile of the PA (An) and PA (Evo) strains used in the study obtained using Kirby-Bauer’s disc diffusion method.

| Antibiotic class | Antibiotic | Antibiotic susceptibility profile |  |
| --- | --- | --- | --- |
|  |  | PA (An) | PA (Evo) |
| Carbapenem | Ertapenem | S | S |
|  | Imipenem | S | S |
|  | Meropenem | S | S |
| Glycopeptide | Teicoplanin<sup>#</sup> | R | S |
|  | Vancomycin | S | S |
| Cephalosporin | Cefotaxime | S | S |
|  | Cefepime | S | S |
| Macrolide | Azithromycin | S | S |
|  | Erythromycin | R | R |
| Penicillin | Methicillin | R | R |
|  | Penicillin G<sup>#</sup> | R | S |
|  | Ampicillin<sup>#</sup> | R | S |
|  | Piperacillin | S | S |
| Tetracycline | Tetracycline | S | S |
|  | Doxycycline | I | S |
|  | Tigecycline<sup>#</sup> | R | S |
| Aminoglycoside | Streptomycin | S | S |
|  | Kanamycin<sup>#</sup> | R | I |
|  | Gentamycin | S | S |
|  | Tobramycin | S | S |
|  | Amikacin | S | S |
| Quinolone | Moxifloxacin | S | S |
|  | Lomefloxacin | S | S |
|  | Ofloxacin | S | S |
|  | Norfloxacin | S | S |
|  | Ciprofloxacin | S | S |
|  | Nalidixic acid | R | R |
| Sulfonamide | Co-trimoxazole<sup>#</sup> | R | S |
|  | Trimethoprim | R | R |
| Rifampicin | Rifampicin<sup>#</sup> | R | S |
| Chloramphenicol | Chloramphenicol | R | R |
| Lincosamine | Clindamycin | R | R |
| Monobactam | Aztreonam | S | S |
| Nitrofurantoin | Nitrofurantoin | R | R |

S- susceptible, R- resistant, I- intermediate
<sup>#</sup> Antibiotics showing collateral sensitivity

### Biofilm formation

Biofilm formation is a significant contributor towards the development of colistin resistance. In this study, the biofilm formation ability of the strains was compared. Three independent experiments indicated that the PA (Evo) strain consistently produced more biofilm than its ancestral counterpart PA (An). Estimation of the biofilm formation ability indicated that the PA (Evo) produced 27.11 % more biofilm than the PA (An) (Fig. 2E).

### Pyocyanin production

In the comparative analysis of pyocyanin production at 24 and 48 hours, it was observed that both the strains produced pyocyanin, but the parental strain produced more pigment (Fig. 2F). After 24 hours of incubation, the PA (An) produced 6.59 µg/ml pyocyanin. In contrast, the PA (Evo) produced 1.72 µg/ml. The same trend was also observed after 48 hours of incubation, where PA (An) and PA (Evo) strains produced 9.47 µg/ml and 3.94 µg/ml of pyocyanin respectively. From this study, it can be concluded that the evolved strain is compromised in its pyocyanin production ability.

### Comparison of the TCS modules

Colistin resistance in *P. aeruginosa* is primarily mediated by the outer membrane remodelling directed primarily by the TCSs PhoPQ and PmrAB. We asked if any alterations in the genes encoding PhoPQ and PmrAB TCS systems are present in the PA (Evo) strains that may be responsible for this strain’s high resistance to colistin. The *phoQ, phoP, pmrB*, and *pmrA* genes were amplified from both the PA (An) and PA (Evo) strains and compared at the nucleotide level. The multiple sequence alignment for the genes *phoQ, phoP*, and *pmrA* did not show any sequence variation between the PA (An) and PA (Evo) strains (data not shown). Interestingly, thymine to guanine substitution at position 407 of the *pmrB* CDS led to a missense mutation from valine to glycine at position 136 (V136G) of the PmrB histidine kinase was detected (Fig. 3A). The mutation is located in the predicted periplasmic domain of the PmrB sensor kinase (Fig. 3B)

**Fig. 3.**
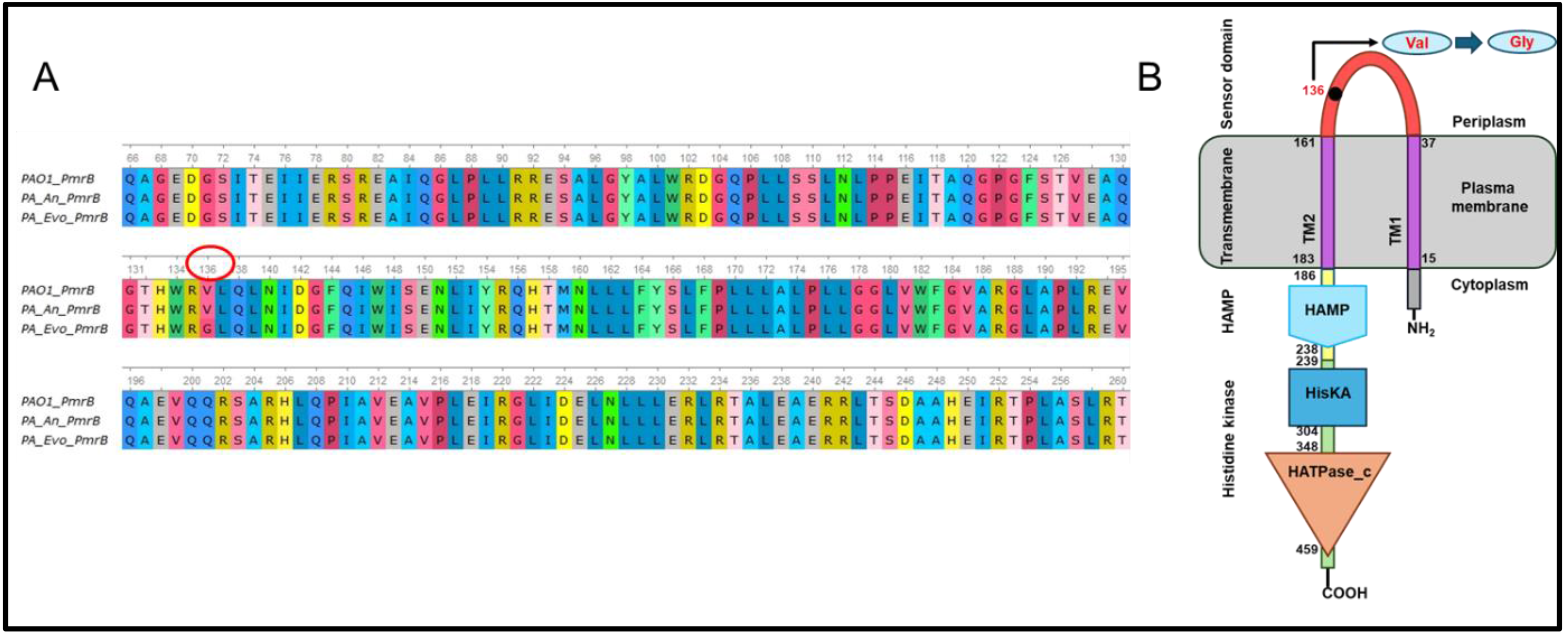
**(A)** Multiple sequence (amino acid) alignment of the PmrB sensor kinase obtained from the PA (An) and the PA (Evo) strains used in the study. The sequence is compared with the PAO1 strain of *P. aeruginosa* as a control. The amino acid substitution (V to G) at position 136 is shown using a red circle. A part of the alignment is shown for clarity. **(B)** The domain architecture of the PmrB sensor kinase was obtained from the SMART database. The amino acid 136 is present in the periplasmic sensor domain and is highlighted in the schematic diagram.

Though various amino acid substitutions have previously been observed in the PmrB kinase in *P. aeruginosa* (Huang et al., 2020; Disney-McKeethen et al., 2023), the V136G detected in this study has not been reported earlier. To understand the impact of the novel amino acid substitution on the functioning of the PmrB kinase, we did the protein variant effect analyzer (PROVEAN) analysis using the provean webserver. The analysis predicted that the V136G substitution would be deleterious with a provean score of -4.087 (cutoff -2.5). However, whether the biological function of the PmrB kinase is affected by the V136G substitution *in vivo* remains to be seen.

## Discussion

*P. aeruginosa* possesses a remarkable ability to develop resistance to frontline drugs in a short period (Laborda et al., 2021; Rossi et al., 2021). Emerging resistance to the last option antibiotic colistin is a significant challenge in the clinic among nosocomial infectious pathogens including *P. aeruginosa* (Padhy et al., 2024). Adaptive laboratory evolution (ALE) experiments are increasingly being used to mimic the growth of pathogens exposed to increasing concentrations of antibiotics in the clinic and the data obtained reflects on the possible evolutionary trajectory to drug resistance (Lenski, 2017; Van Den Bergh et al., 2018; Vinchhi et al., 2023). In the present study, an ALE experiment was employed to evolve a *P. aeruginosa* colistin-sensitive strain PA (An) to highly resistant one PA (Evo), which could grow in the presence of 130 µg/ml of colistin, with gradual passage at increasing concentrations of colistin (Fig. 1). Interestingly, in a span of 106 generations, a 32-fold increase in the colistin MIC was observed in the PA (Evo) strain (Table 2; Fig. 2D). Previous evolution experiment using a morbidostat culture have shown a 10-fold increase in colistin resistance in 10 days and 100-fold in 20 days indicating the rapid rate of resistance emergence (Dößelmann et al., 2017). In this context, we found a 32-fold increase in the colistin MIC in 16 days of passage.

In general, the evolution of drug resistance is often accompanied by a fitness trade-off that may result in an increase or decrease in fitness in different environmental conditions (Barbosa et al., 2017; Andersson et al., 2020; Baquero et al., 2021). This study also found that the PA (Evo) strain formed smaller colonies on LB agar and took a longer time to appear indicating a lower growth rate (Fig. 2A). On comparing the growth characteristics of the PA (An) and PA (Evo) strains in the liquid medium, the latter was found to have slower growth rate, increased lag and generation time, and decreased yield (Fig. 2B, 2C; Table 2). It is interesting to note that though the strain was adapted for 106 generations in the presence of colistin, the evolved strain has lost some of its growth characteristics (fitness trade-off) even in the absence of the selection.

A remarkable attribute of *P. aeruginosa* is the ability to form biofilms at the site of infection which are refractile to antibiotic effect (Ciofu and Tolker-Nielsen, 2019; Sindeldecker and Stoodley, 2021). On comparing the biofilm-forming ability of the strains, we found that the PA (Evo) strain produced more biofilm than the ancestral strain (Fig. 2E). Though, in this study, we have not determined the level of colistin resistance in the biofilm structure, based on the magnitude of resistance observed among the planktonic cells, it can be speculated that the biofilm would be highly resistant to colistin.

Pyocyanin produced by the *P. aeruginosa* strain is a pigment largely involved in virulence and has antibacterial properties (Costa et al., 2017; Zhu et al., 2019). In the present study, the PA (Evo) strain was found to produce significantly less amount of pyocyanin in comparison to its ancestor (Fig. 2F). Interestingly, previous studies have underscored the role of pyocyanin in the formation of biofilm in *P. aeruginosa* (Ramos et al., 2010), indicating the loss of pyocyanin production leading to an overall decrease in biofilm formation. However, in the present study, the PA (Evo) strain produced more biofilm while the pyocyanin production was reduced. This contrasting observation could be due to strain differences and the experimental conditions used. As observed in the present study, the final growth yield of the PA (Evo) strain was less in comparison to its ancestral counterpart often reaching the stationary phase at a lower cell density (Fig. 2B). It may also be possible that the decreasing production of pyocyanin is a result of fitness trade-off with the acquisition of colistin resistance.

Another important aspect of the evolutionary acquisition of drug resistance is that it is often accompanied by collateral sensitivity and cross-resistance (Barbosa et al., 2017; Vanderwoude et al., 2024). Collateral sensitivity is when a strain while acquiring resistance to a drug simultaneously becomes sensitive to other drugs. In contrast, cross-resistance is the development of resistance to a second drug while acquiring resistance to the first one. In this study, we found that the PA (Evo) strain also developed resistance to polymyxin B. As both colistin and polymyxin B belong to the same group of lipopeptide antibiotics a common mechanism would be responsible for the cross-resistance, which is also evident in previous studies (Lázár et al., 2013; Barbosa et al., 2017). Interestingly, the colistin-resistant PA (Evo) strain developed collateral sensitivity to several antibiotics such as penicillin G, ampicillin, teicoplanin, streptomycin, kanamycin, co-trimoxazole, and rifampicin, as determined using the antibiotic susceptibility test. As these antibiotics belong to different classes with distinct modes of action, it is pertinent to understand how sensitivity to all of them developed simultaneously. The PA (Evo) strain might be modified in its cell envelope to restrict the entry of colistin, in turn becoming more accessible by other antibiotics that may gain entry to the cell more efficiently. Studies have shown that sensitivity to β-lactams and aminoglycosides in *P. aeruginosa* is often associated with PmrB-mediated membrane remodelling (Lázár et al., 2013; Baym et al., 2016; Barbosa et al., 2017). As the PA (Evo) strain was found to have a distinct mutation in the *pmrB*, it is plausible that the membrane remodelling impacting the colistin resistance could have made the strain more sensitive to the β-lactams and aminoglycosides. However, this aspect requires experimental validation.

Colistin resistance is predominantly developed by the modification of the bacterial outer membrane mediated by the two-component signalling systems (Padhy et al., 2024). As reported previously in *P. aeruginosa*, PhoPQ and PmrAB are the principal TCSs involved in outer membrane remodelling in response to membrane-acting drugs (polymyxins) and other stresses targeting the bacterial membrane (Miller et al., 2011; Moskowitz et al., 2012). Specifically, various mutations in the sensor kinase and the response regulator components of the TCS either make them defunct or constitutively active regulating the downstream operons (*arnBCADTEF* and *eptA*) involved in cell membrane remodelling. To determine if any mutations accumulated during the 106 generations of adaptive evolution in the PA (Evo) strain, the PhoPQ and PmrAB modules were amplified and sequenced. Interestingly, the PhoPQ and the PmrA response regulator remained unchanged in the PA (Evo), whereas a single substitution mutation was detected in the sensor kinase PmrB. This valine-to-glycine substitution at the amino acid position 136 (V136G) is in the periplasmic sensor domain of PmrB. Several mutations responsible for the development of colistin resistance have been identified in the *pmrB* gene (Dößelmann et al., 2017; Huang et al., 2020; Disney-McKeethen et al., 2023). Moreover, it has been observed that mutations in the PmrAB TCS tend to appear early in the adaptation process followed by other genes in the regulon (Dößelmann et al., 2017). Interestingly, though both PmrAB and PhoPQ are involved in the development of colistin resistance in the experimental evolution studies, data have shown that *pmrB* mutations are rather prevalent in the unstructured cultures as used in the current study, whereas *phoQ* mutations were more prevalent in the structured environment of the microfluidic devices (Disney-McKeethen et al., 2023). Therefore, it is significant to understand the molecular intricacies or the signals that primarily activate these TCSs from an infection point of view. The V136G missense mutation in the PmrB has been predicted to be deleterious by the PROVEAN analysis. However, whether this mutation has any role in the emergence of colistin resistance in the strain remains to be determined. Moreover, if any other mutations emerged in the PA (Evo) strain during evolution need to be ascertained by genome analysis.

## Conclusion

The study using an adaptive laboratory evolution approach developed a *P. aeruginosa* strain highly resistant to colistin. In a span of 106 generations, the colistin resistance was increased by 32-fold as determined by the MIC data. Moreover, the evolved strains showed clear fitness loss in the form of growth characteristics. While the evolved strain showed more biofilm formation, it also developed collateral sensitivity to several antibiotics tested. Interestingly, a novel amino acid substitution (V136G) in the sensor kinase PmrB was detected in the evolved strain. Though PmrB mutations have been implicated in the development of colistin resistance, whether the V136G mutation detected in the present study is involved in the increased resistance to colistin remains to be determined.

## Acknowledgements

This work was supported by the funds received from the “Science and Engineering Research Board (SERB)”, Govt. of India (grant no. SRG/2019/001703). Indira Padhy is a recipient of the “Biju Patnaik Research Fellowship (BPRF)” from the Science and Technology Department, Govt. of Odisha. We thank Namrata Biswal and Truptirani Chand for their assistance in some of the experiments.

## Author contributions

Designed the study: SSM & IP; Performed the experiments: IP & SKD; Data analysis: IP, SKD, & SSM; Manuscript writing: IP & SSM. All authors agreed to the submitted version.

## Conflict-of-interest statement

The authors declare no conflict of interest.

## Notes

### Competing Interest Statement

The authors have declared no competing interest.

